# Uncovering the Genomic Landscape of *Mycobacterium bovis* in Wales

**DOI:** 10.1101/2025.10.07.680915

**Authors:** Amy J E Healey, Cate L Williams, Nicholas J Dimonaco, Terry Galloway, Richard J Ellis, Eleftheria Palkopoulou, Amanda J Gibson, R Glyn Hewinson, Jessica C A Friedersdorff

**Affiliations:** Centre of Excellence for Bovine Tuberculosis, Department of Life Sciences, Aberystwyth University, Aberystwyth, Wales; Institute for Global Food Security, School of Biological Sciences, Queen’s University Belfast, Belfast, Northen Ireland; Animal and Plant Health Agency Carmarthen Field Services, Ty Merlin, Heol Glasdwr, Parc Pensarn, Carmarthen, SA31 2NF; Surveillance and Laboratory Services Department, Animal and Plant Health Agency Weybridge, New Haw, Surrey. KT15 3NB; Department of Bacteriology, Animal and Plant Health Agency Weybridge, New Haw, Surrey. KT15 3NB

**Keywords:** Bovine tuberculosis, MTBC, SNPs, molecular epidemiology, population structure, Wales

## Abstract

Bovine tuberculosis (bTB), caused by the bacterium *Mycobacterium bovis*, is one of the most pressing animal health issues in Wales today. It negatively impacts cattle health, affects profitability and trade, and can decimate years of genetic improvement towards desirable production traits. It also places a significant burden on the health and well-being of affected farming families. Eradication of bTB requires the understanding of local transmission pathways to target effective disease control interventions. Traditional genotyping methods, such as spoligotyping and VNTR analysis, which have previously been used in Wales to understand the epidemiology of M. bovis, have lacked the discriminatory power to accurately identify local transmission pathways. Recently, whole-genome sequencing has replaced traditional genotyping methods for all *M. bovis* strains isolated from infected animals in Wales. In this study, we characterised the genomic diversity of *M. bovis* across Wales by analysing the genome sequence of all 379 *M. bovis* isolates obtained from culture-positive animals in Wales in 2021. Analyses uncovered three main clusters that are geographically distinct. A further three clusters containing fewer isolates were also geographically separated, two of which had particularly large SNP distances compared to the majority of other Welsh isolates, suggesting independent introductions of *M. bovis* strains that are not endemic to Wales. Fine-scale and epidemiologically relevant genetic structuring was identified within the six main clusters, indicating region-specific evolution, which can drive local disease dynamics. Finally, A number of SNPs in coding genes were identified that have the potential for important advantageous physiological consequences that may impact host-pathogen interactions and necessitate further investigation.

## 1. Introduction

*Mycobacterium bovis* is the pathogen that causes bovine tuberculosis (bTB) in cattle and zoonotic TB infections in humans (Brites et al., 2018). Globally, *M. bovis* is found to be maintained in several different wildlife reservoirs beyond cattle (Smith et al., 2006). Bovine tuberculosis is an economically significant disease and is endemic amongst animal hosts in the UK. Whilst Scotland has achieved bTB-free status, cases in England and Wales remain high; by the end of September 2022, 2,145 herds were declared not officially bTB free with 3,645 new herd incidents in 2023 (Spencer, 2011; TB Hub, 2023). In England and Wales, the estimated mean total consequential cost per break-down is £23,636 (Barnes et al., 2023), but the social and psychological implications for farmers and herd managers are currently impossible to quantify (Barnes et al., 2020).

*M. bovis* belongs to the *Mycobacterium tuberculosis* complex (MTBC), alongside at least six other species that cause similar respiratory symptoms in a variety of hosts, including humans. MTBC is a clonal complex, thought to have originated from a single common ancestor with members collectively sharing 99.95% genetic identity (Blouin et al., 2012; Galagan, 2014; Guimaraes & Zimpel, 2020). Horizontal gene transfer (HGT) and recombination events are rare within MTBC; therefore, due to the clonal nature of the MTBC, almost all variation is observed through single nucleotide polymorphisms (SNPs) and insertion/deletion events (indels), most of which are located in repeat regions (Kao et al., 2016; Guimaraes & Zimpel, 2020). *M. bovis* belongs to the animal-adapted lineage of the MTBC and is categorised into clade A4 with *M. caprae* and all *M. bovis* BCG vaccine strains (Brites et al., 2018). As a species, *M. bovis* has been further organised into definitive clonal complexes that are separated geographically. The true number of clonal complexes is yet to be determined, but it is widely accepted that there are at least four: African 1 and 2 (AF1 & 2) and European 1 and 2 (EU1 and EU2) (Smith et al., 2003, 2012). The complex EU1 accounts for almost all animal infections in the UK but has also been identified in several different countries worldwide (Smith et al., 2011; Zimpel et al., 2020).

Whole genome sequencing (WGS) has superseded traditional typing methods to characterise *M. bovis* isolates and for molecular epidemiological studies (Sandhu et al., 2025). Studies using WGS in conjunction with traditional typing methods such as spoligotyping and variable number of tandem repeats (VNTR) have found that SNP-based phylogenies from WGS data separate samples geographically and demonstrate strong local dynamics (Reis et al., 2021). As such, using SNP patterns, in combination with existing molecular methods or independently, has been shown to provide greater epidemiologically relevant granularity than traditional methods. For example, genotypic differences between *M. bovis* isolates from outbreaks of bTB from across the USA (Joshi et al., 2012) and UK (Sandhu et al., 2025) were resolved using SNP patterns, where clades defined by SNP data were represented by more than one spoligotype/VNTR profile, demonstrating the ability of SNP-based phylogeny to provide higher resolution than traditional genotyping methods.

The localisation of SNPs across the genome may also have the potential for functional impact. Previous analysis of *M. bovis* isolates using WGS revealed that of all SNPS occurring within virulence genes, 33% of these SNPs occurred in genes associated with lipid transport and metabolism (Perea et al., 2021). Perhaps not surprisingly, given the importance of the lipid cell wall and its role in virulence and interactions with host immune cells, these genes are also subject to heavy selection pressure by the diagnostic skin test, specifically the single intradermal comparative cervical skin test (SICCT) (Kurz & Rivas-Santiago, 2020). It is therefore essential to provide a functional context for the SNPs used to genotype *M. bovis* strains.

Recently, the Animal Plant Health Agency (APHA) replaced genotyping of field isolates by spoligotyping and VNTR analysis (Waller et al., 2022) with the use of WGS for *M. bovis* isolates cultured from lymph nodes of reactor animals in England and Wales and have recommended a shift to a SNP-based classification system to replace the genotype scheme that has been used previously (Sandhu et al., 2025). In this study, we complement the annual APHA bTB report from 2021, which contains descriptive epidemiology reports on bTB in Wales by analysing the genome sequence of all *M. bovis* isolates obtained in Wales in 2021 to elucidate the population structure of *M. bovis* in Wales and assess the distribution and potential effects of SNPs in these isolates.

## 2. Methods and Materials

### 2.1. Sample preparation and sequencing

As part of the bTB statutory surveillance programme, all *M. bovis* culture-positive isolates collected during routine surveillance in 2021 were characterised by WGS. In multiple-reactor incidents only a sample of reactors are cultured, and generally no more than three animals with lesions are cultured per incident. Usually, only one culture-positive sample is collected from each incident for WGS. Samples were inoculated on modified 7H11 slopes (Gallagher and Horwill, 1977) for 6-12 weeks, and a single colony was harvested for molecular characterisation. These procedures were performed in Containment Level 3 (CL3) laboratories located at the Surveillance and Laboratory Services Department at APHA. Paired-end libraries were constructed directly from heat-inactivated material using the Nextera XT DNA Library Preparation Kit (Illumina, Cambridge, UK), and sequencing was performed at the Central Unit for Sequencing and PCR (CUSP) at APHA on an Illumina MiSeq or NextSeq 500/550 instrument, generating 150bp paired-end reads.

### 2.2. Bioinformatic and phylogenetic analysis

Analysis was carried out using the bespoke Aberystwyth *M. bovis* pipeline (AMBoP) (v1.6.1) which was created in house for this analysis. The code can be found on GitHub (github.com/Aber-TB/AMBoP). AMBoP is a pipeline that takes in raw, unprocessed reads, systematically uses tools to identify SNPs, and creates a variety of outputs, including trees and information on functional effects, whilst giving the user control over each parameter and step during the analysis. In brief, read quality assessments were undertaken before and after trimming using FASTQC (v0.11.9; Andrews, 2010) and multiqc (v1.13; Ewels et al., 2016). Sliding window quality trimming was used in Trimmomatic (v0.39, Bolger et al., 2014). For trimming, a 10 bp window size and quality of 20 was used with a minimum read length of 36, and trimming adapters for various Illumina library preps were removed with 2 seed mismatches, 30 palindrome clip threshold and 10 simple clip thresholds (as per APHA parameters and as described in the GitHub repository btb-seq from APHA-CSU^1^)

Paired trimmed reads were then aligned to a reference genome using BWA (v 0.7.17; Li & Durbin, 2009) and unmapped, non-primary and supplementary alignments were removed using SAMtools (v1.16.1; Danecek et al., 2021) as well as sorting, indexing and removing duplicates. For this study, the reference used was the *Mycobacterium bovis* AF2122/97 genome assembly (GenBank LT708304.1). Samples were retained if >90% of the sites in the reference genome had a coverage depth of >=10. Furthermore, read taxonomy was assigned using Kraken (v2.1.2; Wood et al., 2019). If <70% of the raw reads from a sample were assigned ‘Mycobacterium’ in the taxonomy name but the sample passed the coverage and depth filters, then the taxonomy of reads that aligned to the reference genome were checked to determine the level of contamination (to evaluate whether non-Mycobacterial reads were aligned to the reference genome). This serves as a rescue step for samples that contain both Mycobacterial reads and contaminants, ensuring that only Mycobacterial reads are retained.

SNPs were called using BCFtools (v1.16; Danecek et al., 2021) and resulting variant call files (vcf) were filtered; mapping quality (MQ) >=30, depth of reads at site of alternative allele (DP) >= 10, number of supporting forward and reverse reads was >=1 each, and proportion of reads needed to support an alternative allele was >=0.95. For tree building, where a SNP was within 10 bp of other called variants, all were removed, as these loci can be associated with repetitive regions (Gardy et al., 2011) and/or are likely in linkage disequilibrium and can be indicative of non-neutrality in that genomic region. For returning functional information about SNPs these variants were retained. Sites within known hypervariable or repeat regions were masked and SNPs in these regions excluded, for example, those named as PE or PPE genes, labelled as direct repeats, invert repeats, repeat regions or mobile genetic elements in the annotation file for the genome available from GenBank (GFF3 available from NCBI Reference Sequence: NC_000962.3).

Consensus FASTA files were created for each sample using the reference genome with alternative alleles in positions of predicted SNPs using BCFtools. SNPs present across all samples were located with snp-sites (v2.5.1; Page et al., 2016), creating a pseudo-genome (of polymorphic sites concatenated together) for each sample. SnpEff (v5.1, Cingolani et al., 2012) was used to predict the impact of SNPs and identify those located in functional genes. To get additional functional information, such as Clusters of Orthologous Genes (COGs), eggNOG-mapper online (v5.0; Huerta-Cepas et al., 2019; Cantalapierdra et al., 2021) was used with default settings to annotate the protein-coding genes from the reference genome, obtained from NCBI.

Maximum-likelihood phylogenies were constructed using the pseudo-genomes with RAxML (v8.2.12; Stamatakis, 2014) with fast bootstraps (100) and GTRCAT model, as well as IQ-tree (v2.2.0.3; Minh et al., 2020) using constant sites and with ultrafast bootstrapping (1000 replicates) (Hoang et al, 2018) and the best-fit model (TVM+F+I+I+R7) as identified by ModelFinder (Kalyaanamoorthy et al, 2017) Trees were rooted using the reference strain AF2122/87. The resulting phylogeny was visualised and annotated in iTol (v7.2; Letunic & Bork, 2024). Spoligotypes were obtained using SpoTyping (v2.1, Xia et al., 2016) and the octal codes were queried in the *M. bovis* Spoligotype Database (Smith & Upton, 2012).

Analyses to further define population structure were conducted in R v2.4.2 (R Core Team, 2021). Briefly, concatenated SNP alignments were subjected to fast-hierarchical Bayesian Analysis of Population Structure (fast-BAPS; Tonkin-Hill et al., 2019) using a hierarchical Bayesian clustering algorithm (Heller & Ghahramani, 2005) and the optimised ‘symmetric prior’ to determine the optimal number of clusters at multiple levels of hierarchy. FastBAPs analysis was conducted using Fast-BAPs (version 1.0.8) and plotted with ggplot2 (version 3.4.0; Wickham et al., 2016) and ggtree (version 3.4.4; Yu et al., 2017). Genetic distances between samples were explored using both the number of SNP differences and model-based genetic distances, calculated using ape (version 5.6-2, Paradis et al., 2004), and visualised using pheatmap (version 1.0.12) and ggtree in R v2.4.2 (R Core Team, 2021). Scripts for downstream data analysis and creation of figures can be found on GitHub^2^.

### 2.3. Functional Analysis of SNPs

Using the Eggnog annotations for the reference genome, SNPs that occurred in coding regions were grouped from samples within clusters and plotted using ggplot2, ggpubr (version 0.5.0, Kassambara, 2022) and reshape2 (version 1.4.4; Wickham, 2007) in R v2.4.2 (R Core Team, 2021).

Counts of SNPs in genes were normalised using the gene length to account for the inherent likelihood that longer genes have more SNPs. Counts of SNPs in COG groups were normalised using the total number of genes in that COG group. Genes containing SNPs were also cross referenced with lists of known virulence genes (Hauer et al., 2019), and gene essentiality (Gibson et al. 2021).

## 3. Results

A total of 477 samples were available for this study from Wales in 2021, each consisting of paired-end reads sequenced from a putative M. bovis sample recovered from predominantly bovine hosts, as well as from badgers, fallow deer, and a cat. Of the 477, 98 samples failed the coverage filtering step, where <90% of the reference genome was covered by a minimum of 10x read coverage. There were a further six samples that passed the coverage filtering, but Kraken analysis revealed that <70% of the raw reads were annotated as belonging to the *Mycobacterium* genus (i.e. failed filtering for contamination/indiscriminate sequence identity). However, the read depth was adequate to reach the threshold of reference genome coverage, and of these reads that mapped to the reference, <0.3% were reads annotated by Kraken as something other than *Mycobacterium.* Therefore, these samples were not filtered out and were included in the analysis, resulting in a total of 379 samples. A total of 2,047 SNPs were identified across all 379 samples across Wales.

### 3.1. Phylogenetic Analysis using SNPs Reveals *M. bovis* Population Structure of Wales

After filtering of SNPs within 10 bp of each other, a subset of 1,971 SNPs (of a total of 2,047) were used to create phylogenies and for further downstream analyses. Maximum likelihood (ML) phylogenies (Figure 1) partitioned the dataset into six distinct clusters, supported by bootstrap (BS) values of 100 for each partition and by fast-BAPS Bayesian clustering (innermost ring in Figure 1; Supplementary Figure 2). Spoligotypes were determined for the 379 samples (middle ring in Figure 1) and aligned with BAPS clustering. Some geographic localisation of clusters was evident (Figure 2), particularly for clusters 3, 5 and 6. Cluster 3 was found primarily in samples from southeast Wales, cluster 5 was found exclusively in southwest Wales, and cluster 6 was the dominant genetic lineage in north Wales. Although almost all samples were obtained from cattle, some were also obtained from wildlife, which did not cluster into distinct lineages and were distributed amorphously throughout the tree.

**Figure 1:**
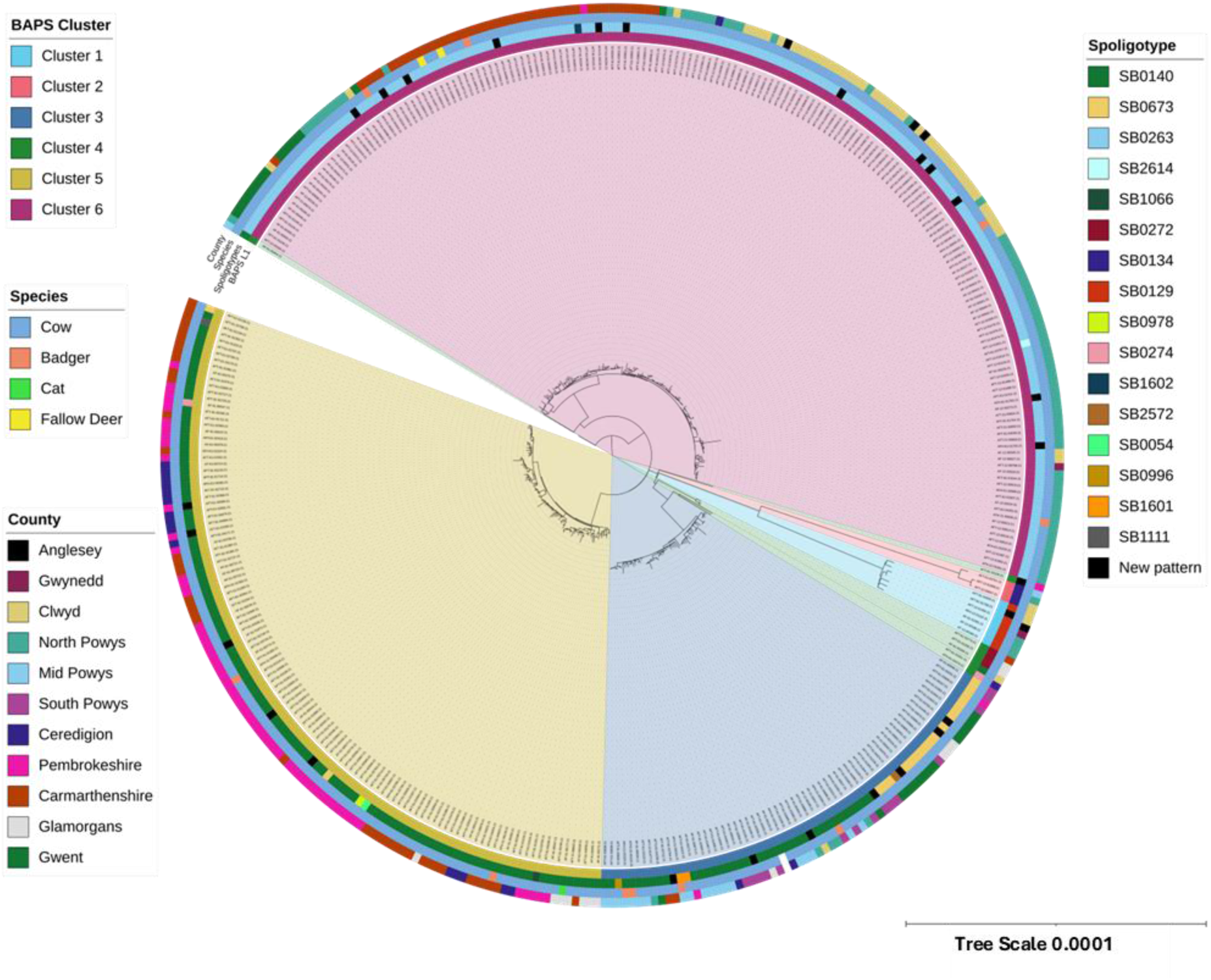
Maximum likelihood phylogenetic tree of *M. bovis* inferred from 1,971 SNPs from 379 isolates across Wales. The reference strain AF2122/87 was used to root the tree. The phylogeny is annotated with coloured concentric rings; from inside out, these represent the level one BAPs cluster, the spoligotype, the county the sample was obtained in, and species the sample was obtained from. Trees were constructed using IQTree and the TVM+F+I+I+R7 model.

**Figure 2:**
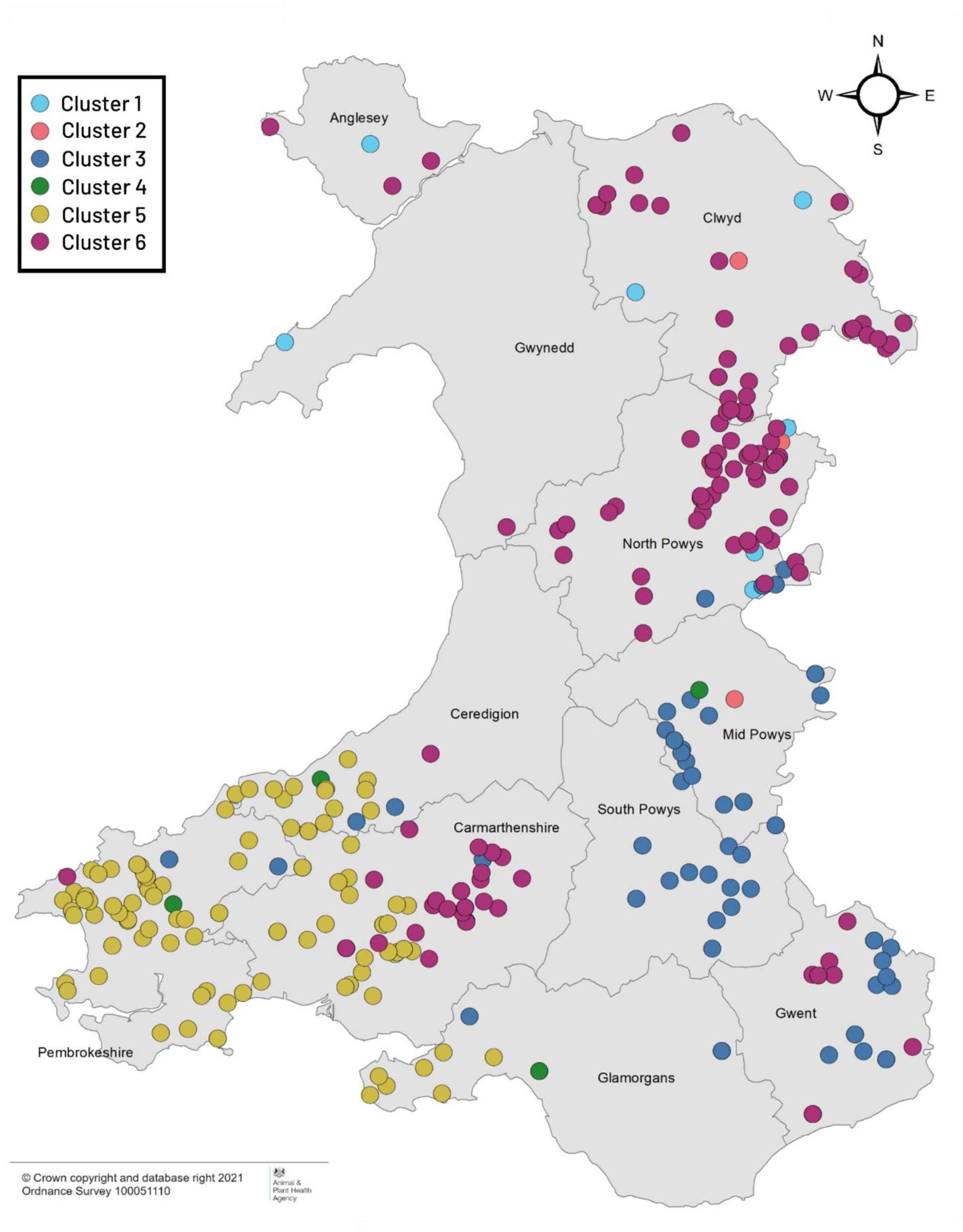
Map of sample locations across Wales. Sampling points are coloured to represent the membership of each sample to each of the six genetic clusters identified by BAPS analysis, where light blue is cluster 1, pink is cluster 2, dark blue is cluster 3, green is cluster 4, yellow is cluster 5 and purple is cluster 6. Note that the county Powys has been split into three for ease of discussion.

The average SNP distance (Supplementary Tables 1 and 2, Supplementary Figure 3) between clusters ranged between 122 (stdev= 4.51) and 516 (stdev= 8.07) SNPs, with clusters 1 and 2 the most distinct, differing from all other clusters by 406-475 and 474-505 SNPs respectively. All the other clusters (3-6) differed from each other by fewer SNPs (122-160). The diversity within clusters (Supplementary Table 3) varied, with the average SNP difference within cluster 4 (mean = 129, stdev=44.22) far greater than within all other clusters, where average SNP distance ranged between 17 and 23. This indicates that cluster 4 likely comprises multiple highly distinct genetic subgroups.

Within the six main clusters, further within clade clustering of isolates was observed using the fastBAPS algorithm (Supplementary Figures 1 and 2) and supported by Maximum Likelihood (ML) BS values of 100 for the nodes separating each monophyletic sub-cluster in the ML-derived phylogeny. When the locations of these sub-clusters were mapped (Figure 3), geographically localised subclustering was resolved within clusters 3, 5 and 6. There were two divergent groups within cluster 3, the first group was restricted to the far south-east of Wales, and the second group was found primarily in the more northern samples in eastern mid Wales, Cluster 5 was partitioned into four sub-clusters, which showed clear spatial localisation within southwest Wales. Cluster 6 also comprised four sub-clusters, which were largely localised to Carmarthenshire (sub-cluster A), Gwent (sub-cluster B), north-east Wales and Anglesey (sub-cluster C) and north Powys (sub-cluster D). Additional shallow population genomic structuring was resolved within sub-clusters.

**Figure 3:**
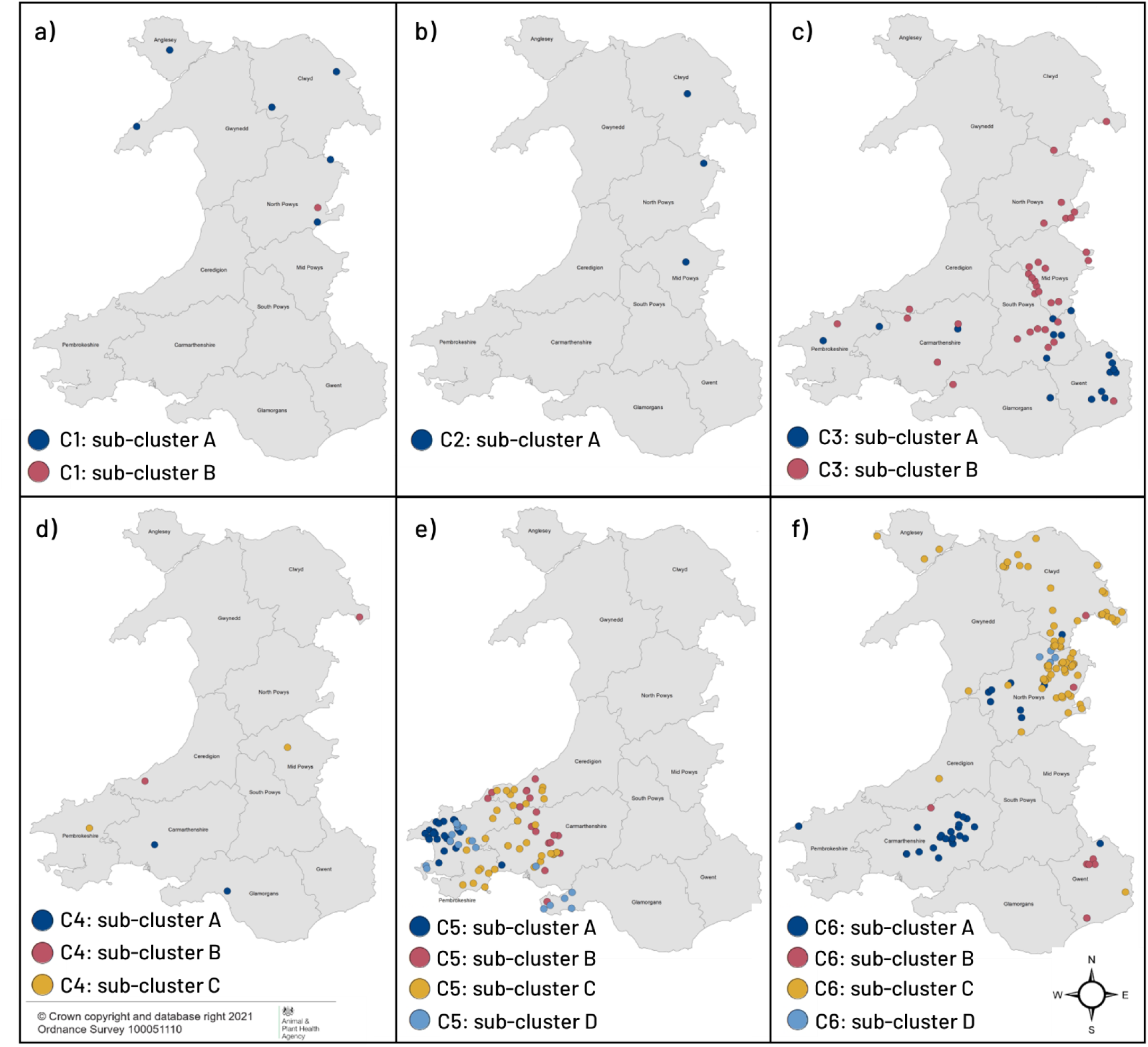
Maps of the distribution of each of the clusters identified by BAPS analysis. a) Cluster 1, b) Cluster 2, c) Cluster 3, d) Cluster 4, e) Cluster 5, f) Cluster 6. Sample points are colour coded based on membership to each of the within cluster (C1-C6) subclusters defined by level 2 BAPS analysis.

### 3.2. SNP Effects on Function

Of the 2,047 variants identified across all samples, 1,770 SNPs occurred in protein coding regions, 271 in non-coding regions and a further six occurred in ribosomal RNA regions. There were three SNPs that occurred in the overlapping region between two genes (both genes are counted separately in the functional analysis). There were 651 SNPs that resulted in synonymous mutations, and 1,120 in non-synonymous (1,104 missense and 16 nonsense) mutations, giving a ratio of 0.58 synonymous to non-synonymous SNPs. The spread of SNPs across different COG functional groups were profiled for each BAPs cluster and compared (Figure 4; Supplementary Table 4). Generally, no single cluster showed a bias towards an accumulation of SNPs in any one functional group. Instead, the frequency of SNPs amongst different functional groups roughly correlates with the proportion of the genome these functions occupy. After normalisation for gene length, the highest proportion of SNPs were found in COG category Secondary Structure (Q) for Cluster 1, Inorganic ion transport and metabolism (P) for Cluster 2, Cell Cycle Control and Mitosis (D) for Cluster 3, Replication and Repair (L) for Cluster 4, Cell Wall/Membrane/Envelope Biogenesis (M) for Cluster 5, and Transcription (K) for Cluster 6.

**Figure 4:**
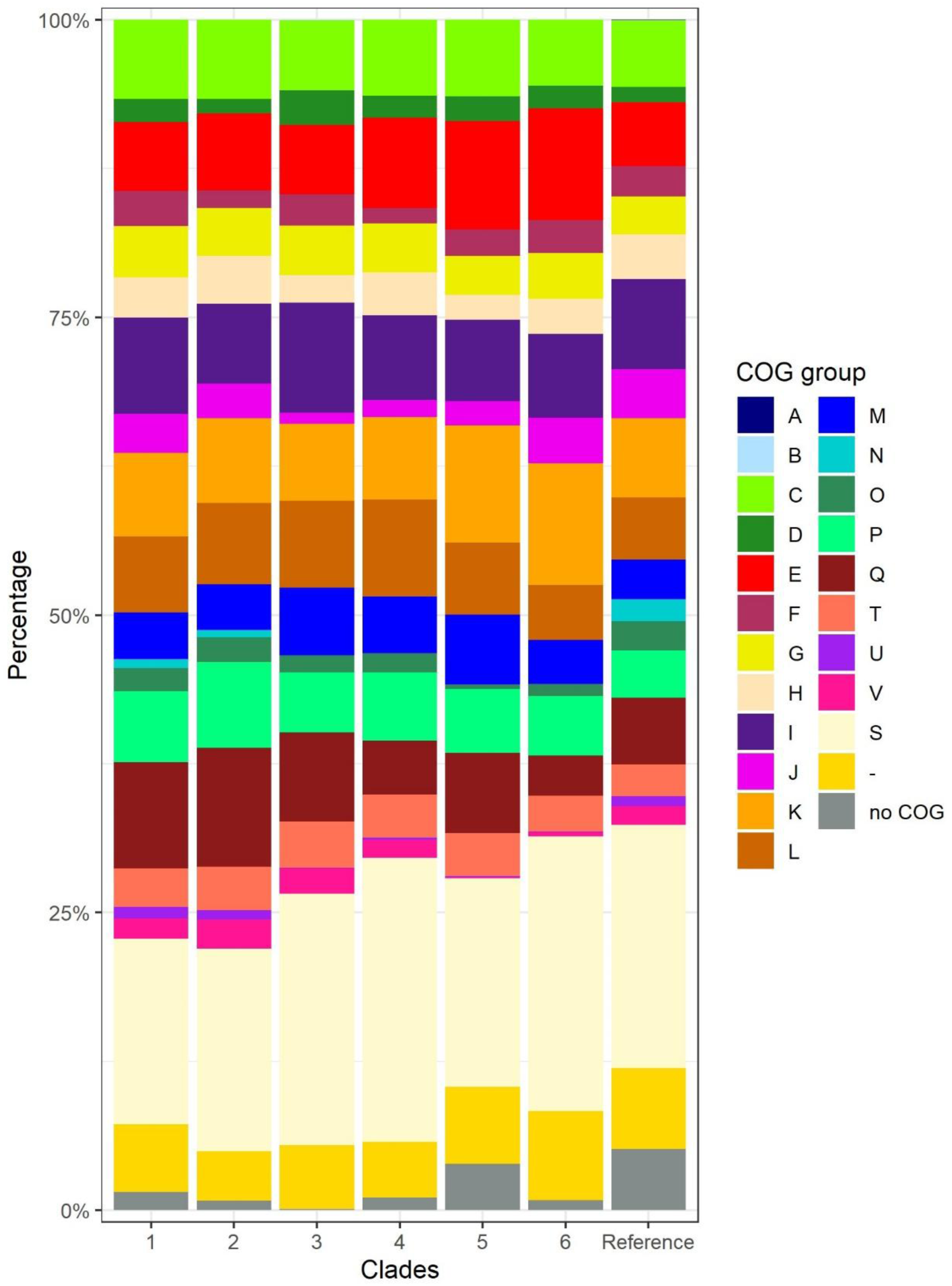
Stacked bar graph with the functional breakdown of SNPs in genes that belong to different COG groups for the six clusters. The proportion of genes belonging to the different COG categories in the reference genome is given for context.

Based on *M. bovis* essentiality allocations made by Gibson et al. (2021), the present study found a total of 1,354 SNPs in 958 non-essential (NE) genes, 225 SNPs in 166 essential genes, 48 SNPs across 38 genes that confer a growth advantage and 143 SNPs across 87 genes that confer a growth disadvantage. Of the essential genes with SNPs, those genes involved in intermediary metabolism and respiration harboured the highest percentage of SNPs at 34%. Around 60% of SNPs in essential genes were missense mutations, the remainder were silent. There was one nonsense mutation in a conserved protein (Mb0504c; orthologous to Rv0493c (Kapopoulou et al, 2011) that confers a growth advantage that was present in seven isolates. In addition, there was one missense mutation found in 10 samples that was marked as high impact and results in a loss of a stop codon in a probable ABC transporter protein (Mb1013; orthologous to 5’ end of Rv0987), which confers a growth disadvantage. Out of 181 virulence genes listed previously (Hauer et al., 2019), 58 genes contained a total of 114 SNPs. Of these, three SNPs were reported as high impact as the SNP causes a stop gain in genes *plcD* (Mb1784c), *ribA1* (Mb1975, ortholog Rv1940), and *lipR* (Mb3111; ortholog Rv3084). A further 67 SNPs were missense mutations in 41 virulence genes, and 44 were silent mutations in 23 virulence genes.

No SNPs were found in any samples in the genes encoding the antigens ESAT-6 (*esxA*, Mb3905), CFP-10 (*esxB*, Mb3904), and Rv3615c (*espC*, Mb3645c), nor were any SNPs found in genes encoding the sero-diagnostic antigens MPB70 (Mb2900) and MPB83 (Mb2898). The gene *pks12* (Mb2074c), a polyketide synthase enzyme, was found to contain the greatest number of SNPs. However, at 12,456 bp, this is also the longest gene in the genome. When normalised for gene length, the gene with the highest frequency of SNP mutations was a conserved hypothetical protein (Mb0772) with 0.0122 SNPs/BP. There were 351 genes containing more than one SNP.

There were 25 SNPs tagged as high impact by SnpEff, which caused start loss (n=4), stop gain (n=16) or stop loss (n=6). All but two of the high-impact SNPs were found in non-essential genes. Of these, three were found in virulence genes (described above), and 11 were private mutations (i.e. they were only identified in 1 of the 379 samples analysed), and the distribution of these were spread relatively evenly across the phylogeny. One SNP which causes the loss of a stop codon in a probable FadB3a protein (Mb1742; ortholog Rv1715) occurred in 378 samples. In *M. bovis* this protein is split into Mb1742 and Mb1743 but exists as a single protein in *M. tuberculosis*. In 46 isolates, a stop codon was gained in a transcriptional regulator of the AcrR family (Mb0848c; ortholog Rv0825c). These 46 isolates all belong to Cluster 5 (Figure 1) and show some geographical localisation; found in Carmarthenshire, south Ceredigion, and Pembrokeshire, an emergent hotspot known as the High TB west Area of Wales. Furthermore, this SNP is unique to all members of Cluster 5 subcluster C.

There are 10 isolates together from Cluster 1 (n=7) and Cluster 2 (n=3) that all contained the same four high-impact missense SNPs, which were unique to these two clusters, each predicted to result in the loss of a stop codon from that gene (Mb0731, Mb1013, Mb1351c, Mb2607). An additional four high-impact SNPs were unique to Cluster 1, three of which result in premature stop codons and one in the loss of a start codon. Another three SNPs were unique to Cluster 2, all of which resulted in a premature stop codon (Supplementary Figure 1). These clusters appeared to form phylogenetically distinct lineages and thus were also tested for the presence of a SNP in *guaA* (Mb3429c) which differentiates lineages in the EU2 clonal complex. None of these isolates, or any others in this study, had this defining SNP (G A) at position 3765573 (Rodriguez-Campos *et al*, 2012). Further identification of these divergent clusters is beyond the scope of this study.

## 4. Discussion

This study utilised WGS data for all *M. bovis* isolates collected across Wales in 2021 from infected cattle (and from badgers, fallow deer and a cat) to define the population structure of *M. bovis* in Wales. These analyses complement the annual bTB epidemiology and surveillance reports published by APHA (e.g. APHA 2020; 2021), providing fine-scale spatial resolution for Welsh *M. bovis* isolates. Furthermore, WGS data was utilised to identify functionally significant SNP mutations throughout the phylogeny that could have advantageous physiological effects.

### 4.1 Population Structure of Welsh *Mycobacterium bovis*

WGS data provided considerable resolution for evaluating population structure of epidemiological relevance, and sheds light on the physical and evolutionary factors that shape the distribution and abundance of *M. bovis* in Wales. The genome-wide SNP patterns partitioned the *M. bovis* isolates in the 2021 dataset into six distinct clusters, of which three groups were dominant: Cluster 3 (17.1% of all Welsh samples), Cluster 5 (31.1%), and Cluster 6 (47.1%). In addition to these three main genetic lineages, a further three less common genetic clusters were identified (Clusters 1, 2 & 4). Both Clusters 1 and 2 were highly genetically divergent from the rest of the Welsh *M. bovis* population. These genetic differences, as well as phylogenetic partitioning, suggest that Clusters 1 and 2 have been diversifying along their own evolutionary trajectories for a significant amount of time and originate from different ancestral lineages than the *M. bovis* reference strain, AF2122/97. The *M. bovis* strain AF2122/97 was derived from a virulent case of bTB in Great Britain (GB) in 1997, and is assigned to the spoligotype SB0140 and the EU1 clade, which is common within the UK. Isolates in clusters 1 and 2 were assigned to SB0129 and SB0134 respectively, which have been shown to be more closely related to French strains (Garcia Pelayo et al., 2009), with SB0134 spoligotypes labelled as a yet undefined clonal complex (Loiseaeu et al., 2020; van Tonder et al., 2021), supporting the hypothesis of an alternate ancestor to the reference strain. The small number of isolates identified within Clusters 1 and 2 suggests that these are uncommon genotypes in Wales.

There was a clear geographic basis to the distribution of the three dominant genetic types across Wales; Cluster 3 was the dominant genetic lineage in mid and southeast Wales, Cluster 5 was found only in southwest Wales, and Cluster 6 was found primarily in northeast Wales, as well as in two localised and spatially aggregated clusters in Carmarthenshire and Gwent. These results are consistent with the WGS clade home ranges for Wales in 2021 reported recently by Seery et al. (2024) using analysis performed by APHA. In this publication, the authors reported six WGS clade home ranges within Wales. The major clades identified were the APHA clade B6-11, equivalent to cluster 6, which was the most abundant clade, accounting for 45.9% of the total number of isolates sequenced in 2021. This finding aligns with the results of Sandhu et al. (2025), which identified B6-11 as the most prevalent and widespread clade in the UK. The next most abundant clades were B6-14, equivalent to Cluster 5, which accounted for 32.3% of the total isolates and was almost exclusively located in the southwest of Wales (the high TB west management area - APHA 2021) and clade B6-83, equivalent to Cluster 3, accounting for 16.2% of the total and predominantly localised in Gwent and Powys (The high TB east management area-APHA 2021). The home ranges for B6-11 and B6-83 extend across the border of Wales into England (Seery et al., 2024; Sandhu et al. 2025), whilst B6-14 is unique to Wales, with all cases elsewhere attributed directly to cattle movement from south-west Wales (APHA personal communication).

By comparing the genetic structure resolved here with that of previous studies, we can infer how the spatial distribution and abundance of genetic types in Wales have changed over time. In 2003, the most abundant spoligotypes in Dyfed (a former county that spans Ceredigion, Carmarthenshire and Pembrokeshire), Powys and Gwent were reported as part of a broader UK-wide investigation (Smith et al., 2003). The SB0140 spoligotype (which corresponds with Cluster 3 Subcluster B and Cluster 5) was reported as the most abundant spoligotype in Dyfed and Powys, and was present in Gwent, (as well as being the most common spoligotype across the whole of the UK) (Smith et al., 2003), This remains true in the present study, with Cluster 5 the dominant genetic type in Pembrokeshire, and Cluster 3 Subcluster B the dominant type in mid and south Powys. Spoligotype SB0263 (Cluster 6) was previously the second most common in the UK, reported by Smith et al. (2003), and has since become the most abundant in the UK (Sandhu et al. 2025) and is accordingly the most common spoligotype overall in this current study. This is one of the most geographically dispersed clusters, with isolates present in every county except Glamorgan. SB0673 was the third most common spoligotype identified in the UK by Smith et al. (2003), particularly in Gwent, and corresponds to Cluster 3 Subcluster A, which is predominantly found in Gwent and south Powys.

Within the six main clusters, further within clade clustering of isolates was observed, especially for Clusters 3, 5 and 6, where each sub-cluster was localised to a specific region of Wales. The geographical localisation of *M. bovis* lineages across Wales is likely due to clonal expansion of a dominant strain in the area caused either by the spread of a favourable mutation, together with all other genes present in the ancestral cell in which the mutation occurred, or by the invasion of a novel genotype into a geographical region through cattle movement (Smith et al., 2003; van Tonder et al., 2021). Such region-specific evolution and potential adaptation can drive local disease dynamics and outcomes and have been well-documented for *M. tuberculosis* infection in humans (Gagneux & Small, 2007; Caws et al., 2008; Warner & Mizrahi, 2014). The fine-scale granularity of variation resolved here offers the potential to improve our understanding of the origin of infection and associated local risk pathways for the spread of bTB. This knowledge will help target optimal control interventions, which will be the focus of future studies.

### 4.2 The role of cattle movement and wildlife reservoirs in shaping population structure

The distribution and abundance of *M. bovis* isolates in Wales generally reflect the density of cattle herds, with more samples in Pembrokeshire, Carmarthenshire, and north Powys, and fewer occurrences in the central mountainous regions, where cattle density is lower (APHA 2021). A notable exception to this is Anglesey and the Llyn Peninsula, where, despite a high cattle density comparable to the higher TB areas of Pembrokeshire and north Powys, the prevalence of *M. bovis* is low. The density of hosts is recognised as being of critical importance to the transmission of bTB (Allen et al., 2018; Crispell et al., 2019; Rossi et al., 2021), which necessitates careful consideration in terms of the management of bTB; particularly given the trend towards intensification of farming practices over recent decades (Brooks-Pollock & Keeling, 2009) and highlights the importance of a ‘keep it out’ policy for areas of Wales that have a low incidence of *M. bovis* infection and for herds that are free of tuberculosis.

Cattle movement is well recognised as a major driver of *M. bovis* transmission across long distances (Gilbert et al., 2005; Green et al., 2008; Brooks-Pollock et al., 2014; Rossi et al., 2021; Van Tonder et al., 2021), and the phylogenies reported in this study, when interpreted in the context of reported cattle movement data (APHA 2020 report) suggest that cattle movement may have played an important role in introducing many of the genetic clusters present in Wales. For example, those isolates with distinct SNP patterns and large SNP differences from the other clusters may indicate an imported strain. All isolates in Cluster 2 have the spoligotype SB0134, which has been identified in countries worldwide, including South Africa, Spain, Mali, France, and Brazil (Ghavidel et al., 2018). It was also the most prevalent spoligotype in Ethiopia (Mekonnen et al., 2020) and in Algeria, based on samples taken in 2017 (Belakehal et al., 2022), and one of the most prevalent in France, as indicated by samples from 1978 to 2011 (Hauer et al., 2019). Isolates with this spoligotype are distinct from those of the GB-dominant EU1 clonal complex with Mekonnen et al. (2020), suggesting it did not belong to any of the established clonal complexes in GB. However, samples with this spoligotype have been recorded previously in isolates found in Dyfed, Gwent and Powys, as well as in the English counties of Gloucestershire, Hereford and Worcestershire, and Cornwall (Smith et al., 2003) suggesting isolates with this spoligotype may have been imported, though whether this is multiple independent import events into GB or a single introduction followed by population expansion via intra-GB movement requires further investigation.

Within-cluster phylogenetic partitioning provides further evidence for the role of cattle movement in transmitting distinct genetic lineages across Wales, and sheds light on the demographic events that occur post-transmission. Cluster 6 has undergone a recent spatial expansion and proliferation in northern Wales (APHA, 2016, 2017, 2019, 2020, 2021), with an increasing frequency in northeast Wales, as well as extending its range into Anglesey and Gwynedd between 2019 and 2020 (APHA, 2020). Furthermore, intra-clade genetic structure indicates the presence of two founder events (e.g., Chiner-Oms et al., 2022) of this genetic lineage into Carmarthenshire (Cluster 6: Subcluster B) and Gwent (Cluster 6: Sub-cluster A), which were most likely introduced by the movement of infected cattle into these regions from the historical home range of the respective genetic type.

This was then followed by rapid population expansion (e.g., Rossi et al, 2022; van Tonder et al., 2021), which can be associated with efficient transmission amongst cattle in these regions. Further analyses of temporal variation in these genetic sub-clusters across Wales are required to further understand the drivers and dynamics of these localised outbreaks.

In the British Isles, the Eurasian Badger (*Meles meles*) acts as a wildlife reservoir for maintaining and spreading *M. bovis* infection (Corner et al., 2011). In general, when a pathogen co-circulates in multiple systems (such as multiple host species), it will spread more rapidly, as inter- and intra-specific transmission events result in elevated persistence and multiple species-specific population dynamics diversify the transmission pathways (Craft et al., 2008; Haydon et al., 2002). Badgers have low dispersal potential and restricted home ranges (Corner et al., 2011) and are thus recognised more for their role in the maintenance of bTB within a localised geographic area (Crispell et al., 2019; Rossi et al., 2021; van Tonder et al., 2021), rather than transmission over any substantial distance. This is exemplified by van Tonder et al. (2021), who used M. bovis isolates from the randomised badger culling trial in GB to infer transmission between badgers and cattle. The results suggested that the transmission clusters in different parts of southwest England, which are still evident today, were established by long-distance seeding events involving cattle movement, rather than by recrudescence from a long-established wildlife reservoir. The authors also found that clusters are maintained primarily by within-species transmission, with less frequent spill-over from badgers to cattle and vice versa. With only 10 *M. bovis* isolates obtained from badger hosts in this dataset, it is difficult to draw any substantial conclusions. Although the badger data reported here are limited, the isolates do not show host specificity, as demonstrated by a lack of clustering by host species in the phylogenetic tree. Instead, bTB isolates from badgers are most closely related to isolates from sympatric cattle, which is consistent with previous observations reported for an all-Wales Badger Found Dead survey carried out between 2014 and 2016 (Schroeder et al., 2020). Further investigations are required to understand the local drivers of *M. bovis* transmission. Zimpel et al., (2020) analysed genomes from 24 different hosts, including infected domesticated animals such as bovids, camelids, felines and canines, and wild animals such as mustelids, cervids, felines and identified a lack of host-specific clustering, which further supports the hypothesis that *M. bovis* is a ‘generalist’ member of the MTBC complex that infects a wide range of mammalian species.

### 4.3 The functional potential of SNPs in *M. bovis* in Wales

In total, 1,770 SNPs were found in protein-coding genes. A ratio of synonymous to non-synonymous SNPs of 0.58 was calculated for this dataset, which is consistent with those reported for M. bovis isolates representative of French populations (0.56) (Hauer et al., 2019) and USA isolates (0.49) (Joshi et al., 2012). The number of missense mutations was approximately double the number of silent mutations, with very few nonsense mutations. Missense mutations result in an amino acid change, which may affect protein structure and subsequently function, but equally may not affect the protein sequence; as such, it is difficult to predict the impact of such SNPs (Golby et al., 2013). Non-sense mutations, however, are more likely to affect a change in protein structure and thus function, due to the introduction of a premature stop codon. Of particular interest would be premature stops occurring in regulators, which will impact the expression of one or more genes within a regulon.

Such might be the case with the transcriptional regulator gene in the AcrR family (Mb0848c), where a SNP introduces a premature stop codon, and this mutation occurs in all 46 isolates belonging to Cluster 5, Subcluster 3. To assess whether this SNP confers an advantage, temporal data are needed to determine any population expansion associations with a competitive advantage. Additionally*, in vitro* studies are required to determine its phenotypic effects, alongside a combination of *in vitro* and *in vivo* studies to evaluate its immunological impact on the host.

Across the six clusters and the reference genome, no single COG category contained significantly more SNPs than the others. A hypothesis explored here was whether SNPs congregate in genes with similar key functions (such as lipid metabolism) that together improve the success of *M. bovis* isolates and that these advantageous mutations would form a distinct clade. This, however, was not the case here. Overall, the COG category containing the most SNPs was transcription (K) (excluding genes with unknown function), which is also the functional group with the most genes in the reference genome. This has been highlighted by Bigi et al. (2016), who found variations in 20 regulatory proteins in *M. bovis* compared to *M. tuberculosis* and postulated that these may contribute to host specificity and influence behaviour under hypoxic environments (Bigi et al., 2016). This may reflect the generalised enrichment of bacterial genomes with transcription factors, which enables rapid adaptation to complex environmental conditions through the regulation of gene expression (Cole et al., 1998; Cases et al., 2003; Raman et al., 2004). Furthermore, the lack of SNPs in core virulence factors ESAT-6, CFP-10, and EspC highlights their high degree of conservation, as described in other literature (Encinas et al., 2018), and reinforces the utility of these antigens in the diagnosis of bTB using either skin tests or gamma interferon-based diagnostic assays.

The distribution of mutations across all samples revealed that the occurrence rate of SNPs within essential (E) and non-essential (NE) genes is similar, when taken as a proportion of genes in each category (1,301 SNPs were found in 954 NE genes, 224 SNPs in 166 E genes). Furthermore, missense mutations occurred with equal frequency in both E and NE genes, with 808 NE genes and 133 essential genes having missense SNPs. Of the 16 nonsense SNPs, 15 were found in NE genes, one in a gene conferring growth advantage, and none were identified in essential genes.

SNPs were identified in 89 of the 181 virulence genes described previously in *M. bovis* (Hauer et al., 2019). Of interest are those that SnpEff highlighted as high impact due to nonsense mutations. The *plcD* gene (Mb1784c) is unique to *M. bovis* and has no ortholog in *M. tuberculosis* H37Rv due to the deletion of the RvD2 region by the IS6110 insertion; it is a membrane-bound phospholipase C and, in a very small number of *M. bovis* isolates, is also disrupted by transposition of the IS6110 element (Lari et al., 2001; Viana-Niero et al., 2006). Yet, despite the well-known role of phospholipase C proteins in virulence in several intracellular pathogenic bacteria (including *M. tuberculosis*) its precise role in *M. bovis* is yet unknown (Viana-Niero et al., 2006). The gene *ribA1* (Mb1975) is involved in the riboflavin biosynthesis pathway, where *ribA1* and *ribA2* catalyse the first step in the chain reaction breaking down GTP (Long et al., 2010). The introduction of a premature stop codon and effective knock-out of the gene would not impact riboflavin biosynthesis, possibly due to the compensatory activity of ribA2, supported by the classification of *ribA1* as NE in *M. bovis* (Gibson et al., 2021). The *lipR* gene (Mb3111) encodes an esterase that hydrolyses short-chain esters. In *M. tuberculosis* the esterase was identified in inclusion bodies, and mouse models revealed its ability to inhibit interferon-γ (IFN-γ) and interleukin-2 (IL-2) secretion and stimulation of IL-10 (Chun-Xi et al., 2019). The study by Chun-Xi et al. (2019) demonstrates anti-inflammatory activity via the inhibition of pro-inflammatory cytokines and suggests that this protein plays a role in host-pathogen interaction, at least in *M. tuberculosis*. The introduction of a premature stop codon early in the sequence (base 149/927) would likely knock out the gene, a phenomenon that is frequently seen in *M. tuberculo*sis where the gene contains large sequence polymorphisms resulting in complete deletion of *lipR* in clinical isolates from patients with active disease (Sheline et al., 2009). This suggests that the SNP in Mb3111 may not significantly affect virulence or transmission but may alter the expression of some cytokines in the host during infection.

### 4.4 Not all *M. bovis* isolates have split proteins

Cluster 1 and Cluster 2 are distinct from the other clusters observed in this study, as demonstrated by the long tree branches in the phylogeny (406-516 SNP difference between Clusters 1 and 2, and all other clusters). Isolates in these clusters also possess more unique SNPs with the potential to affect functional changes, as identified by SnpEff as high impact. Some of these SNPs appear to cause the loss of stop codons present in the genomes of *M. bovis* AF2122/97 and members of clusters 3, 4, 5 and 6. Orthologous genes in *M. tuberculosis*, (which shares a common ancestor with *M. bovis* (Brosch et al., 2002), encode fully functional proteins in these differential regions. It is the acquisition of a SNP in the evolution of *M. bovis* that results in a premature stop codon and the splitting of these proteins. There are four examples in the present dataset where split proteins are created by the presence of a stop codon in *M. bovis* clusters 3-6. The gene *atsA* (Mb0731; ortholog Rv0711) encodes an arylsulfatase that exists as a single protein in M. tuberculosis H37Rv, whereas in M. bovis, this gene is split into *atsA* and *atsB* (Kapopoulou et al., 2011). For the *M. tuberculosis* H37Rv gene Rv0987 that encodes an ABC transporter, the stop codon in *M. bovis* results in the protein being split into Mb1013 and Mb1014. For *alkA,* a methyltransferase encoded by the singular Rv317c in *M. tuberculosis* H37Rv, the premature stop codon results in *alkAa* (Mb1351c) and *alkAb* (Mb1350c) in *M. bovis*. Finally, a SNP in *M. bovis* causes a frameshift in the single conserved hypothetical protein Rv2577 in *M. tuberculosis* H37Rv, resulting in Mb2607 and Mb2608 (Kapopoulou et al., 2011). The 10 isolates belonging to Clusters 1 and 2 possess the same alleles for these four genes as *M. tuberculosis* H37Rv, suggesting that perhaps not all *M. bovis* isolates possess split proteins. This evidence suggests that isolates belonging to Clusters 1 and 2 may have descended from a different ancestor than *M. bovis* AF2122/97, one that originated before the acquisition of the SNPs causing the split proteins.

### Conclusion

In this study we characterised the genomic diversity of *M. bovis* across Wales by analysing the genome sequence of all 379 *M. bovis* isolates obtained in Wales in 2021. The use of SNPs identified in *M. bovis* isolates provided greater resolution to traditional methods, such as spoligotyping and VNTR analysis, and revealed the population structure of *M. bovis* in Wales. Analyses uncovered three main clusters that are geographically distinct. A further three clusters, each containing fewer isolates, were also geographically separated, with two of these having particularly large SNP distances compared to the majority of other Welsh isolates, suggesting independent introductions of *M. bovis* strains that are not endemic to Wales. WGS analysis provided enhanced resolution and clarified fine-scale population structure, offering important epidemiological insights into the population dynamics of *M. bovis* in Wales at both local and national levels. Finally, a number of SNPs were identified in coding genes that have the potential for significant advantageous physiological consequences, which may impact host-pathogen interactions.

## 7. Author Contributions

AJEH was responsible for formal analysis, investigation, visualisation, interpretation and writing. CLW contributed to data curation, validation, investigation, writing – original draft. NJD contributed to data curation, methodology and software by consulting on AMBoP. TG assisted with data curation, and visualisation with map creation. RJE and EP facilitated data sharing, transfer and publication. GH was responsible for conceptualisation, funding acquisition. JCAF was responsible for data curation, software and methodology through the creation of AMBoP, formal analysis, visualisation and writing. All authors contributed to writing – review and editing.

## 9. Acknowledgments

The authors would like to colleagues in APHA and the Welsh Government for their support and assistance in providing information relevant to this study and would like to thank and acknowledge James Dale, Karen Gover and the Central Sequencing Unit at APHA, Weybridge for technical support and Joe Crispell whose code available on GitHub inspired and informed decisions and code writing.

## 10. Supplementary Material

**Supplementary Figure 1:**
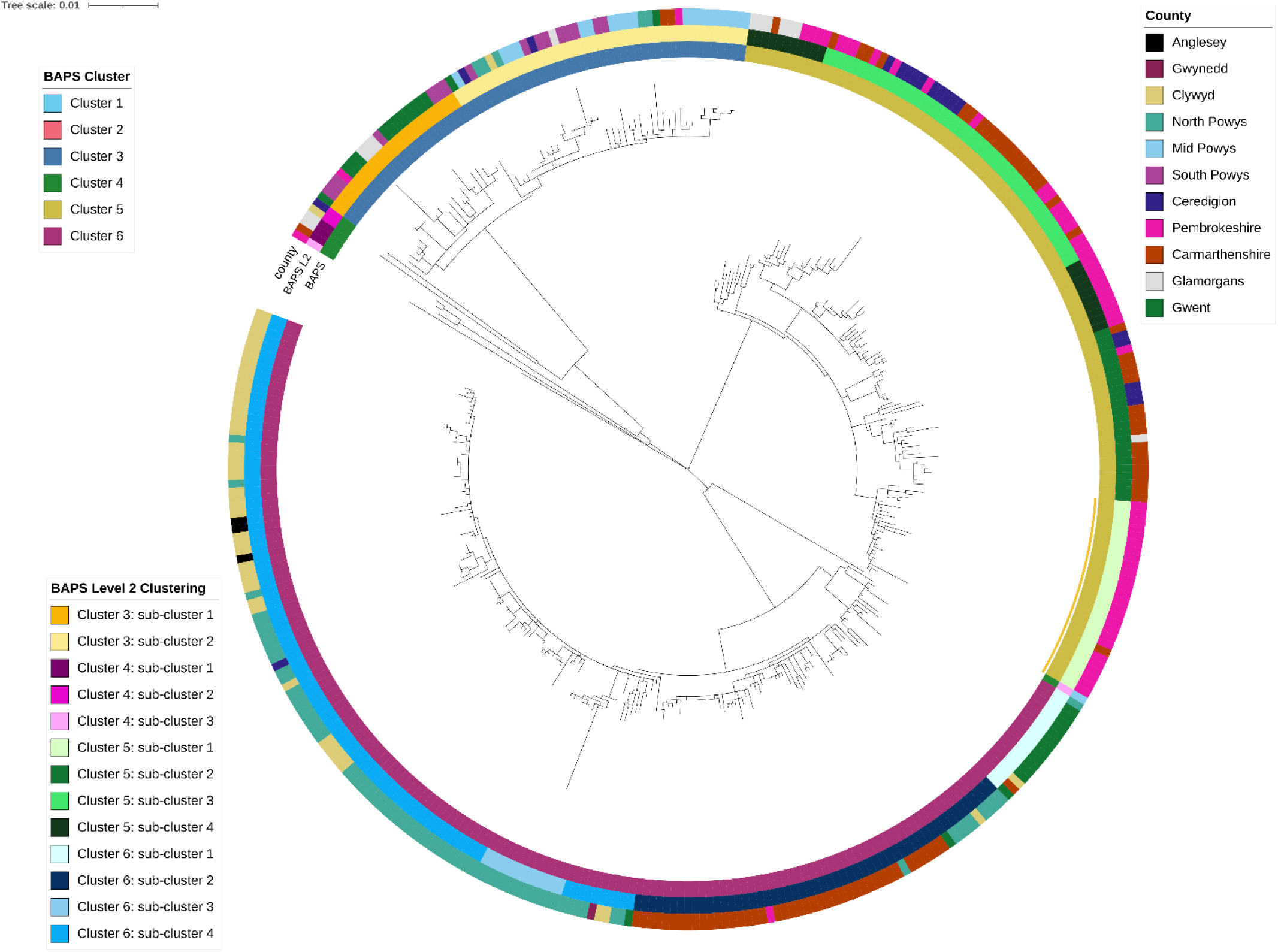
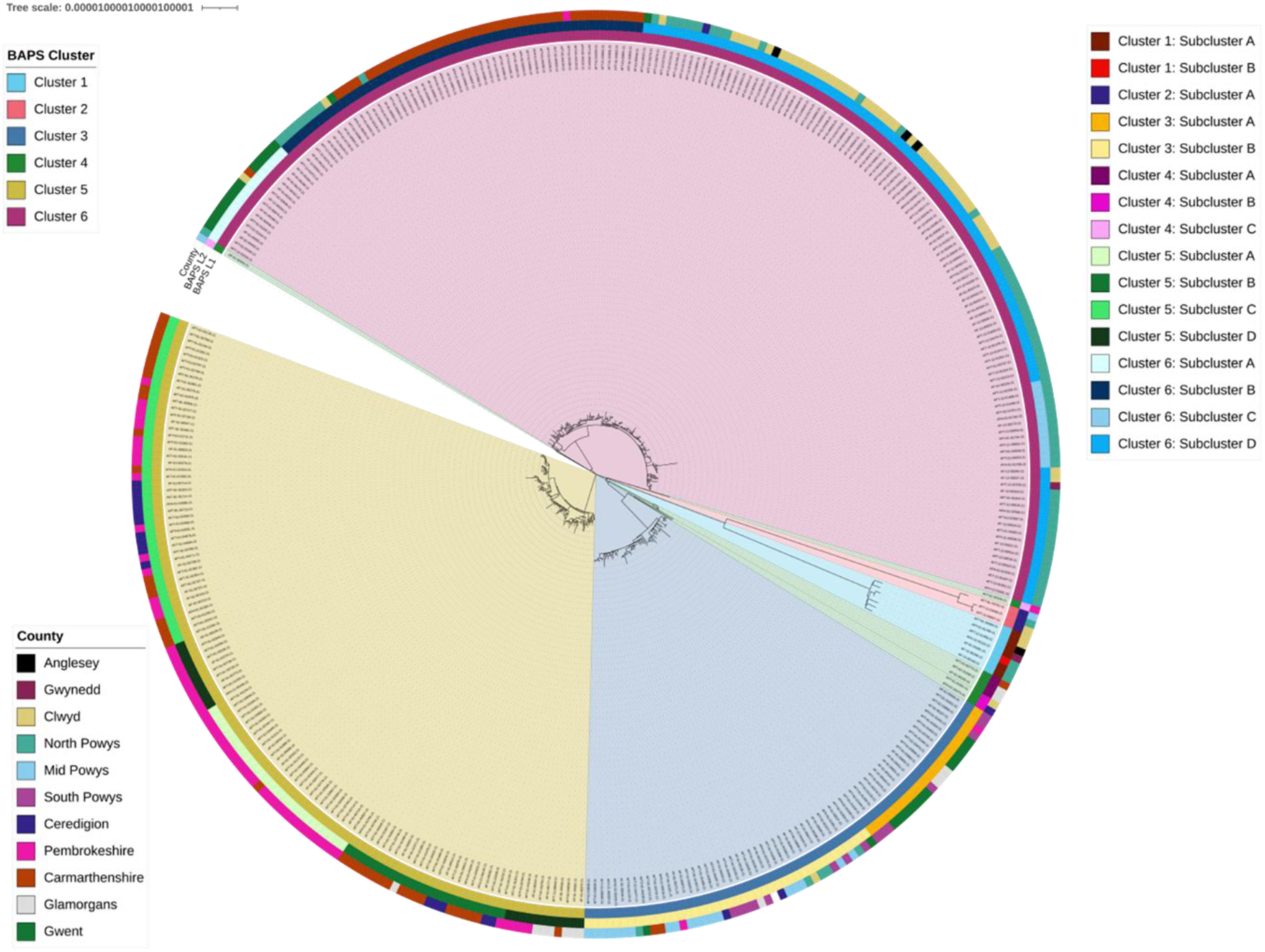
Maximum likelihood phylogenetic tree of *M. bovis* pseudo-genomes, with the divergent clusters (BAPS clusters 1 & 2 in Figure 1) removed. The coloured rings are colour coded to represent: i) the samples that belonged to each Bayesian Analysis of Population Structure (BAPS) cluster, ii) the samples that belonged to each BAPS level 2 (i.e. within group clustering) cluster, and iii) the county in which the sample was isolated from. Tree constructed using IQTree, using model TVM+F+I+I+R7.

**Supplementary Figure 2:**
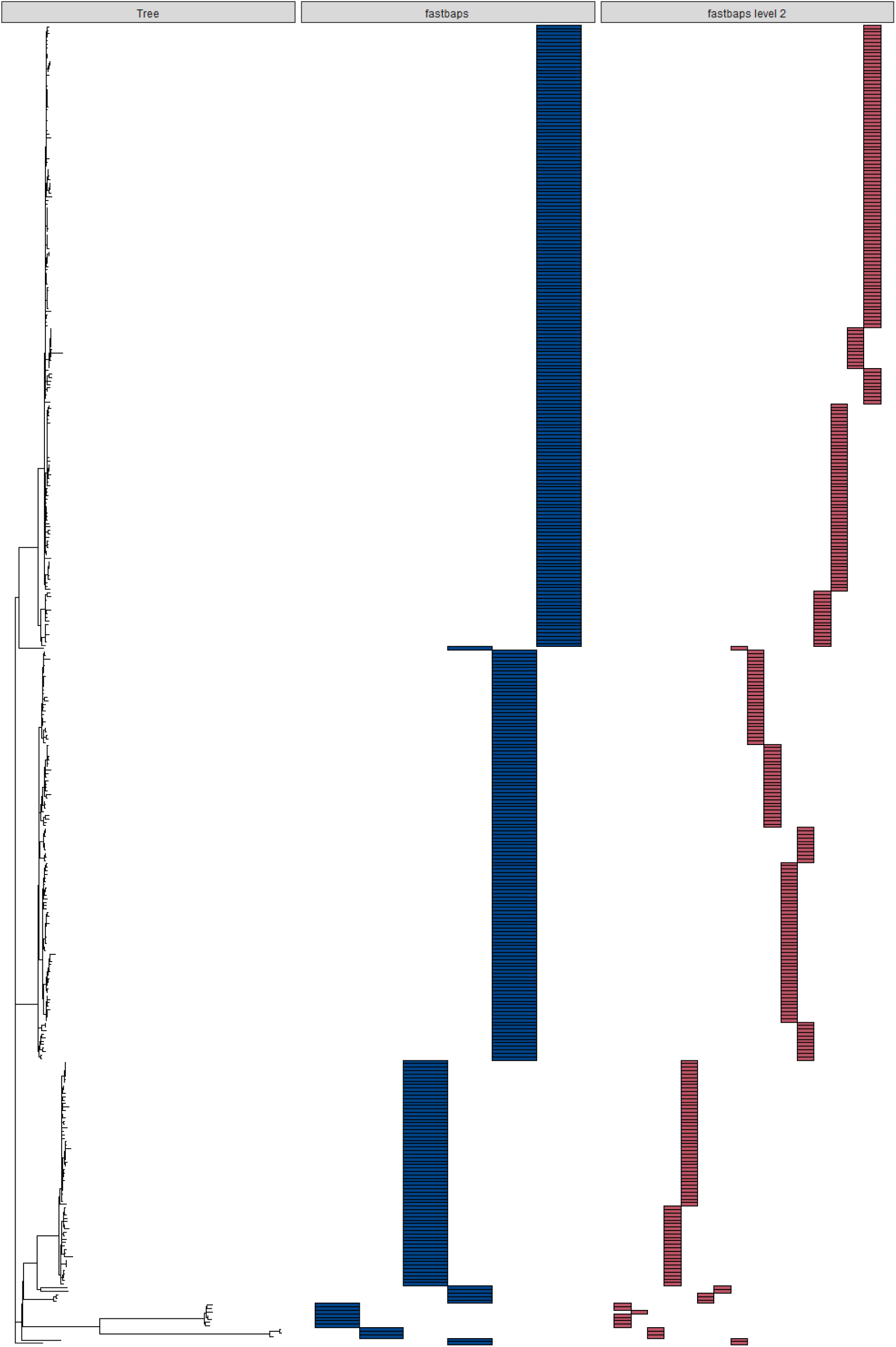
Bayesian clustering of *M. bovis*, inferred from the pseudo-genomes generated from 1,971 SNPs of X *M. bovis* isolates from across Wales, using the fast-BAPS algorithm. Clustering was inferred at multiple levels of hierarchy (level 1 and level 2). The clusters are shown in comparison to the maximum likelihood phylogeny.

**Supplementary Figure 3:**
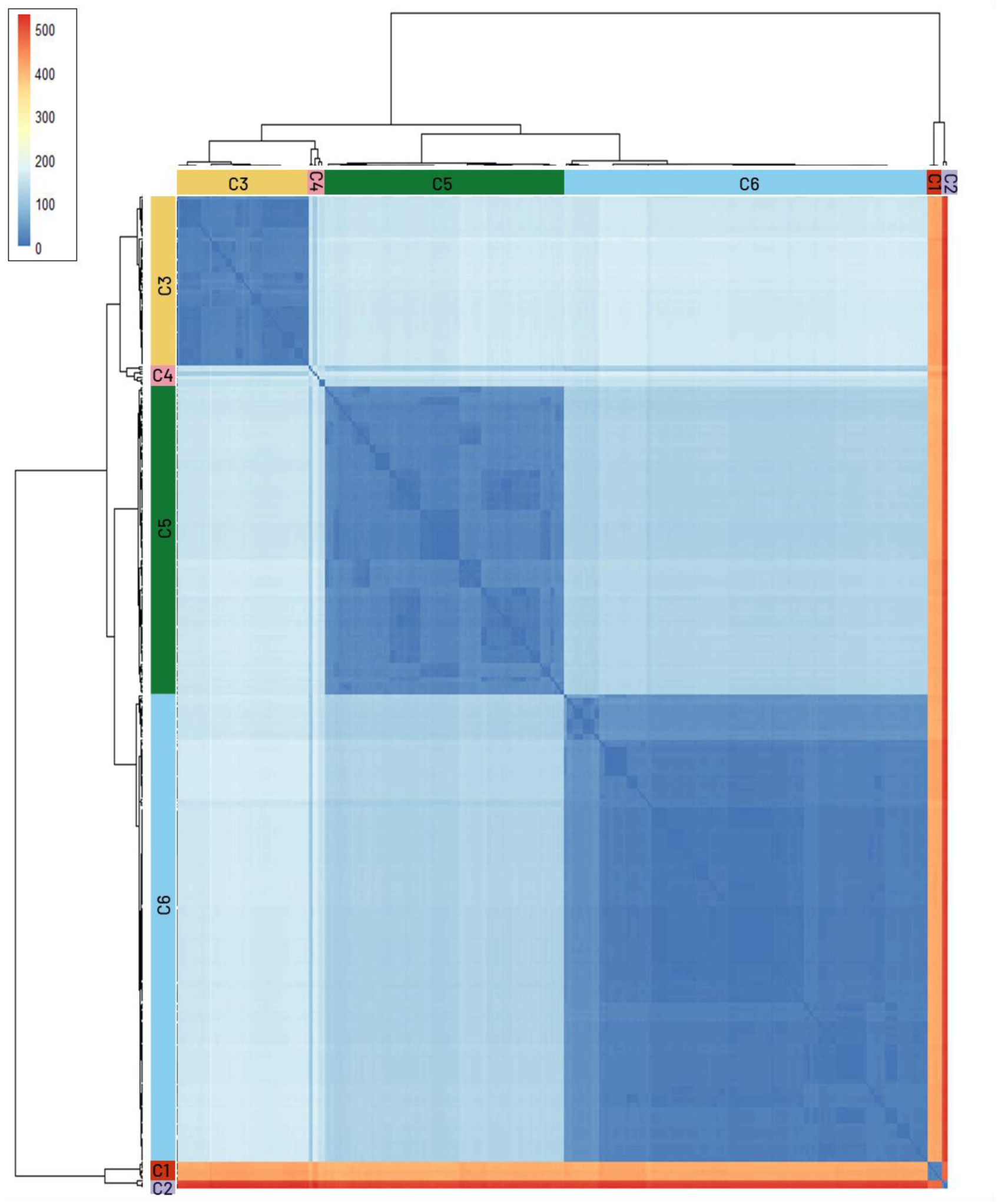
Heatmap of the pairwise SNP distances, calculated as the number of SNPs that differ between *M. bovis* isolates. Dark blue represents low numbers of SNPs that differ between samples and red represents a high number of SNPs. Samples are indicated as their position on the maximum likelihood phylogenetic tree and trees are annotated with the clusters identified by BAPS analysis.

**Supplementary Table 1:**
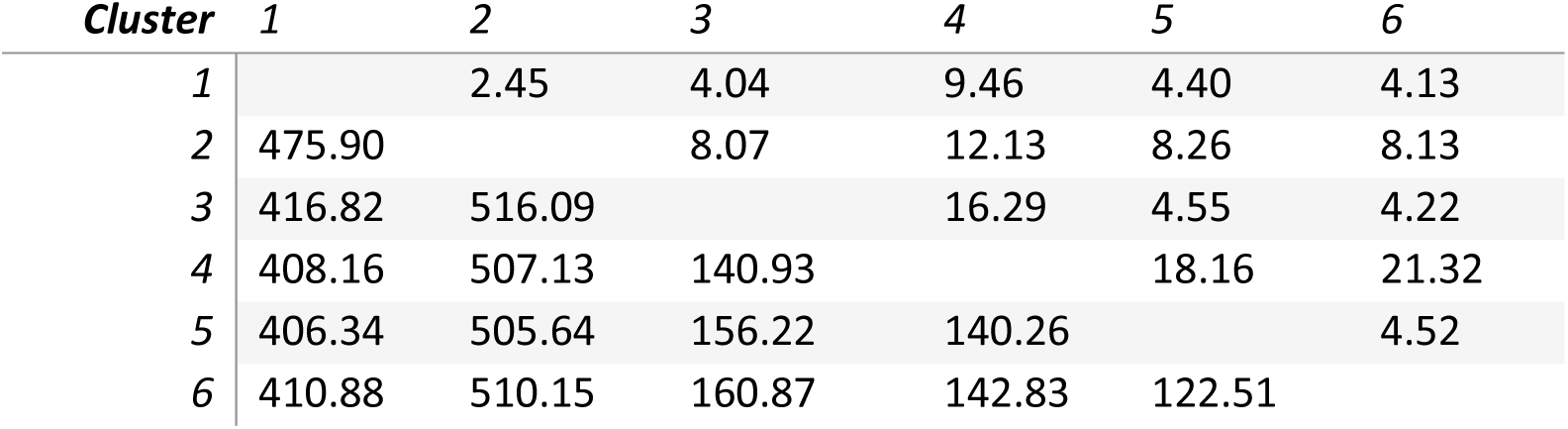
Pairwise SNP differences between each of the clusters identified by fastBAPS analysis. Displayed as the average number of SNP that differ between the members of each cluster below the diagonal and the standard deviation above the diagonal.

| *Cluster* | 1 | 2 | 3 | 4 | 5 | 6 |
| --- | --- | --- | --- | --- | --- | --- |
| 1 |  | 2.45 | 4.04 | 9.46 | 4.40 | 4.13 |
| 2 | 475.90 |  | 8.07 | 12.13 | 8.26 | 8.13 |
| 3 | 416.82 | 516.09 |  | 16.29 | 4.55 | 4.22 |
| 4 | 408.16 | 507.13 | 140.93 |  | 18.16 | 21.32 |
| 5 | 406.34 | 505.64 | 156.22 | 140.26 |  | 4.52 |
| 6 | 410.88 | 510.15 | 160.87 | 142.83 | 122.51 |  |

**Supplementary Table 2:** The minimum-maximum number of SNPs that differ between each of the clusters identified by BAPS analysis.

| *Cluster* | 1 | 2 | 3 | 4 | 5 |
| --- | --- | --- | --- | --- | --- |
| 1 |  |  |  |  |  |
| 2 | 471-480 |  |  |  |  |
| 3 | 407-430 | 500-532 |  |  |  |
| 4 | 394-427 | 487-529 | 109-170 |  |  |
| 5 | 394-419 | 487-521 | 138-175 | 98-172 |  |
| 6 | 398-423 | 491-523 | 142-179 | 95-176 | 106-140 |

**Supplementary Table 3:** Number of SNP differences within the six clusters identified by BAPS analysis. Given as mean number of SNPs that differ between members of that cluster and its standard deviation (stdev) and the minimum and maximum numbers of SNPs that differ between any members of that cluster. N= number of samples assigned to that cluster.

| *cluster* | *N* | *Mean* | *StDev* | *Min* | *Max* |
| --- | --- | --- | --- | --- | --- |
| 1 | 7 | 21.10 | 3.09 | 16 | 26 |
| 2 | 3 | 23.330 | 15.04 | 6 | 33 |
| 3 | 65 | 17.66 | 6.89 | 0 | 42 |
| 4 | 8 | 129.32 | 44.22 | 5 | 166 |
| 5 | 118 | 23.39 | 7.86 | 0 | 44 |
| 6 | 179 | 17.48 | 10.43 | 0 | 64 |

**Supplementary Table 4:** Count of SNPs occurring in COG functional categories for each cluster. The counts of genes within each category in the reference is given for context and to allow for normalisation of the counts.

|  | Cluster |  |  |  |  |  |  |  |  |
| --- | --- | --- | --- | --- | --- | --- | --- | --- | --- |
| COG group | 1 | 2 | 3 | 4 | 5 | 6 | Ref | Total | Total adjusted |
| A | 0 | 0 | 0 | 0 | 0 | 0 | 2 | 0 | 0 |
| B | 0 | 0 | 3 | 0 | 0 | 0 | 1 | 3 | 3 |
| C | 183 | 99 | 543 | 59 | 770 | 1084 | 242 | 2738 | 11.31405 |
| D | 56 | 18 | 269 | 17 | 255 | 371 | 56 | 986 | 17.60714 |
| E | 165 | 96 | 538 | 70 | 1095 | 1833 | 233 | 3797 | 16.29614 |
| F | 85 | 22 | 242 | 12 | 265 | 549 | 109 | 1175 | 10.77982 |
| G | 123 | 60 | 387 | 38 | 395 | 743 | 138 | 1746 | 12.65217 |
| H | 98 | 60 | 211 | 33 | 250 | 583 | 163 | 1235 | 7.576687 |
| I | 231 | 100 | 853 | 66 | 820 | 1374 | 329 | 3444 | 10.46809 |
| J | 95 | 43 | 74 | 13 | 245 | 752 | 178 | 1222 | 6.865169 |
| K | 187 | 102 | 597 | 62 | 1185 | 1993 | 289 | 4126 | 14.27682 |
| L | 169 | 98 | 675 | 71 | 730 | 911 | 225 | 2654 | 11.79556 |
| M | 113 | 57 | 523 | 44 | 706 | 716 | 144 | 2159 | 14.99306 |
| N | 21 | 9 | 0 | 0 | 0 | 1 | 81 | 31 | 0.382716 |
| O | 56 | 31 | 136 | 15 | 43 | 198 | 106 | 479 | 4.518868 |
| P | 171 | 107 | 464 | 53 | 641 | 983 | 173 | 2419 | 13.98266 |
| Q | 152 | 115 | 683 | 42 | 814 | 660 | 241 | 2466 | 10.23237 |
| T | 78 | 50 | 357 | 31 | 436 | 582 | 117 | 1534 | 13.11111 |
| U | 28 | 6 | 5 | 2 | 6 | 12 | 36 | 59 | 1.638889 |
| V | 49 | 36 | 197 | 14 | 8 | 76 | 69 | 380 | 5.507246 |
| S | 447 | 253 | 1947 | 216 | 2102 | 4496 | 885 | 9461 | 10.6904 |
| - | 163 | 62 | 497 | 43 | 776 | 1466 | 294 | 3007 | 10.22789 |
| no COG | 44 | 12 | 9 | 10 | 467 | 163 | 223 | 705 | 3.161435 |

## 11. Data Availability Statement

Whole genome sequencing data was obtained from the APHA under a data transfer agreement. Such data is sensitive and therefore cannot be made public. The scripts for the analysis however are made fully available online.

